# Essential gene analysis in *Acinetobacter baumannii* by high-density transposon mutagenesis and CRISPR interference

**DOI:** 10.1101/2020.09.15.299016

**Authors:** Jinna Bai, Yunfei Dai, Andrew Farinha, Amy Y. Tang, Sapna Syal, German Vargas-Cuebas, Defne Surujon, Ralph R. Isberg, Tim van Opijnen, Edward Geisinger

## Abstract

*Acinetobacter baumannii* is a poorly understood bacterium capable of life-threatening infections in hospitals. Few antibiotics remain effective against this highly resistant pathogen. Developing rationally-designed antimicrobials that can target *A. baumannii* requires improved knowledge of the proteins that carry out essential processes allowing growth of the organism. Unfortunately, studying essential genes has been challenging using traditional techniques, which usually require time-consuming recombination-based genetic manipulations. Here, we performed saturating mutagenesis with dual transposon systems to identify essential genes in *A. baumannii* and we developed a CRISPR-interference (CRISPRi) system for facile analysis of these genes. We show that the CRISPRi system enables efficient transcriptional silencing in *A. baumannii*. Using these tools, we confirmed the essentiality of the novel cell division protein AdvA and discovered a previously uncharacterized AraC-family transcription factor (ACX60_RS03245) that is necessary for growth. In addition, we show that capsule biosynthesis is a conditionally essential process, with mutations in late-acting steps causing toxicity in strain ATCC 17978 that can be bypassed by blocking early-acting steps or activating the BfmRS stress response. These results open new avenues for analysis of essential pathways in *A. baumannii*.

**Importance:** New approaches are urgently needed to control *A. baumannii,* one of the most drug resistant pathogens known. To facilitate the development of novel targets that allow inhibition of the pathogen, we performed a large-scale identification of genes whose products the bacterium needs for growth. We also developed a CRISPR-based gene knockdown tool that operates efficiently in *A. baumannii*, allowing rapid analysis of these essential genes. We used these methods to define multiple processes vital to the bacterium, including a previously uncharacterized gene-regulatory factor and export of a protective polymeric capsule. These tools will enhance our ability to investigate processes critical for the essential biology of this challenging hospital-acquired pathogen.

## Introduction

The Gram-negative bacterium *Acinetobacter baumannii* is among the most difficult to treat pathogens causing diseases in hospitals. *A. baumannii* is an important cause of pneumonia and bloodstream infections and is associated with outbreaks in healthcare environments. The microorganism has rapidly evolved resistance to a wide variety of antimicrobials, leaving few therapeutic options for infected patients. Some strains show resistance to all available antibiotics, including carbapenems and the last-line polymyxins, rendering them exceedingly difficult, if not impossible, to treat (1, 2). There is an urgent need for new antimicrobials that can target the pathogen.

Devising novel strategies to target and attack *A. baumannii* requires that we understand proteins that have essential cell functions. Unfortunately, much remains unknown about fundamental processes in *Acinetobacter*. The genus has diverged from other *Gammaproteobacteria* and lacks sequence orthologs of several important proteins involved in key pathways including cell wall synthesis, cell division, stress responses, and transcriptional regulation (3, 4). Orphan or hypothetical proteins may have evolved to mediate these processes in unique ways in *Acinetobacter*. A number of such proteins were recently identified through functional genomics examination of transposon mutant drug susceptibility phenotypes (5). While this work established functional connections between important biological pathways and uncharacterized proteins, it was designed for analysis of mostly non-essential genes. Phenotypes linked to essential genes, those required by the organism for growth in standard nutrient medium, have yet to be systematically examined in *A. baumannii*, although they will likely provide much-needed insights into pathways enabling growth, stress resistance, and persistence.

Although a number of tools allow genetic analysis in *A. baumannii* (6, 7), studying essential genes is usually an inefficient process. Such genes are typically analyzed by using homologous recombination to introduce an inducible promoter that substitutes for native control elements, allowing conditional expression. Engineering mutants in this fashion, however, is time-consuming and not amenable to scaling for high-throughput analysis. An alternative approach uses the highly transformable, nonpathogenic relative *A. baylyi* to examine terminal phenotypes after direct allelic exchange of essential gene deletions (8, 9). The approach enables a direct view into the consequences of complete gene loss, but such deletions do not allow tunability of gene expression and analogies with *A. baumannii* must be verified.

CRISPR interference (CRISPRi) is a recently described method for easily programmable gene knockdown in a variety of organisms (10). The most widely used CRISPRi systems employ a nuclease-deficient variant of the *Streptococcus pyogenes* Cas9 enzyme (dCas9) that is guided to a target gene by a single guide RNA (sgRNA), a hybrid molecule which incorporates Cas9 scaffolding and DNA targeting sequences (11). Recognition of target DNA depends on both the sgRNA and a protospacer adjacent motif (PAM) within the targeted DNA bordering the sequence being targeted. The bound dCas9-sgRNA complex represses expression of the target gene by sterically hindering transcriptional initiation or elongation (11). A mobilizable CRISPRi system has been developed for use with several pathogens including *A. baumannii* (12), but the reported knockdown efficiency (~10-fold) is likely to be insufficient for analysis of highly expressed genes in the organism.

In this paper, we present a comprehensive set of candidate essential genes in *A. baumannii* identified through high-density transposon mutagenesis, and the development of a CRISPRi system for efficient analysis of these genes. We used these tools to confirm the essentiality of a recently identified orphan protein functioning in cell division, AdvA, as well as a previously uncharacterized protein belonging to the AraC family of transcriptional activators. In addition, we demonstrate the conditional essentiality of late steps in the biosynthesis of capsular polysaccharides. This work identifies new sites of vulnerability in *A. baumannii* and lays the foundation for future large-scale studies of essential gene function in the pathogen.

## Results

### Identification of candidate essential genes in *Acinetobacter baumannii* by Tn-seq

To determine gene essentiality across the *A. baumannii* genome, we performed saturating random transposon mutagenesis in strain ATCC 17978 and identified mutations that allow colonies to form on rich medium. To this end, we used two independent transposition systems which enable random insertional mutagenesis at high efficiency at different sites in the genome: a *Himar1 mariner* transposon system (5), which generates insertions at TA dinucleotides (13, 14), and a Tn*10*-altered target specificity (Tn*10*-ATS) system, which generates insertions effectively at random due to relaxed site specificity (13, 15). We generated two separate, highly saturated mutant libraries from ~550,000 and ~300,000 mutant colonies with each system, respectively (Materials and Methods). We next used massively parallel sequencing of transposon-genome junctions (Tn-seq) to identify the location of transposon insertions within the libraries. Genomic DNA immediately adjacent to transposon insertions was amplified, enumerated by the Illumina platform, and mapped within the chromosome. With the *mariner* library, insertions were mapped to 174,238 unique TA sites out of a total of 269,711 possible sites (Materials and Methods), equivalent to 64.6% of sites hit. With the Tn*10*-ATS library, insertions were mapped to 98,462 unique chromosomal positions. 368,173 mutants with distinct chromosomal transposon insertions were represented across both libraries, equivalent to an average of one insertion approximately every 10 bp.

To identify essential genes, these Tn-seq datasets were analyzed independently using Bayesian/Gumbel methods with the TRANSIT software package (16). These methods identify genes with unusually long, consecutive stretches of potential insertion sites lacking insertions. The probability of long gaps occurring by chance is then calculated by a Bayesian (with *mariner*) or non-Bayesian (with Tn*10*-ATS) analysis of the Gumbel distribution (16, 17). With the *mariner* library, 392 genes passed the posterior probability threshold for essentiality, and 79 genes were called “uncertain” due to having a probability of essentiality not exceeding this threshold (Table S1). With Tn*10*-ATS, 474 genes were called essential. Genes predicted to have high probability of essentiality with both transposition systems represent the most reliable candidates for being essential in the organism, so we analyzed the overlap between hits with both systems, including the essential and uncertain calls with *mariner* and the essential calls with Tn*10*-ATS. This identified 372 genes as hits with both systems (Fig. 1), and we define these 372 genes as the candidate essential gene set in *A. baumannii*. This set of genes corresponded well with candidate essential genes identified in previous Tn-seq studies with the same strain (18) and with the unrelated, MDR strain AB5075 (19), with 72% (267 genes) showing essentiality across all three studies (Fig. S1).

**Fig. 1.**
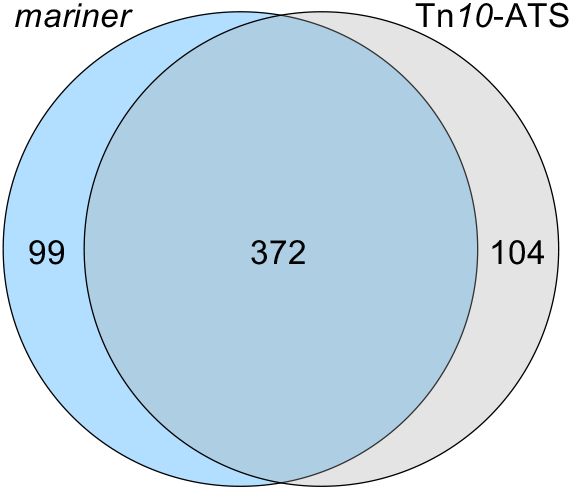
Candidate essential genes in *A. baumannii* identified by Tn-seq with 2 transposon systems. Venn diagram shows candidate essential genes identified in ATCC 17978 by *mariner* transposition (471 genes called essential or uncertain, blue circle) or by the Tn*10*-ATS system (474 genes called essential, gray circle). The intersection of the genes sets had 372 genes, representing the candidate essential gene set using both systems.

### CRISPR interference system for gene knockdown in *A. baumannii*

To facilitate the analysis of the candidate essential genes, we developed a CRISPRi system in *A. baumannii*. The system comprises an anhydrotetracycline (aTc)-inducible *dcas9* inserted at single copy in the chromosomal *attTn7* site downstream of the *glmS* locus (20), and a constitutive sgRNA module via a high-copy plasmid derived from pWH1266 (21) (Fig. 2A). In an initial test of gene knockdown with the CRISPRi system, we targeted the constitutive β-lactamase ADC, levels of which can be determined by measuring rate of hydrolysis of its specific chromogenic substrate nitrocefin (22). Using an sgRNA construct containing 24 nucleotides targeting the non-template (NT) strand of *adc* starting 90 bp downstream of its predicted transcription start site (TSS) (Fig. 2B and Table S2) (23), we observed reduction of β-lactamase levels by almost 2-fold in the absence of *dcas9* induction compared to a non-targeting control plasmid (Fig. 2C). After *dcas9* was induced by 100 ng/ml aTc for 2 hours, β-lactamase synthesis decreased by approximately 30-fold compared to control, approaching the background level seen with deletion of the *adc* gene (Fig. 2C). As expected, *adc* knockdown by CRISPRi increased the susceptibility of *A. baumannii* to ampicillin, a substrate of the ADC enzyme (22, 24). *dcas9* induction completely blocked growth of the strain harboring the *adc*-targeting sgRNA in the presence of a dose of the drug that was sub-inhibitory with control cells, with partial growth inhibition observed in the absence of induction (Fig. 2D). These results show that efficient gene knockdown can be achieved in *A. baumannii*, enabling investigation of gene-phenotype relationships.

**Fig. 2.**
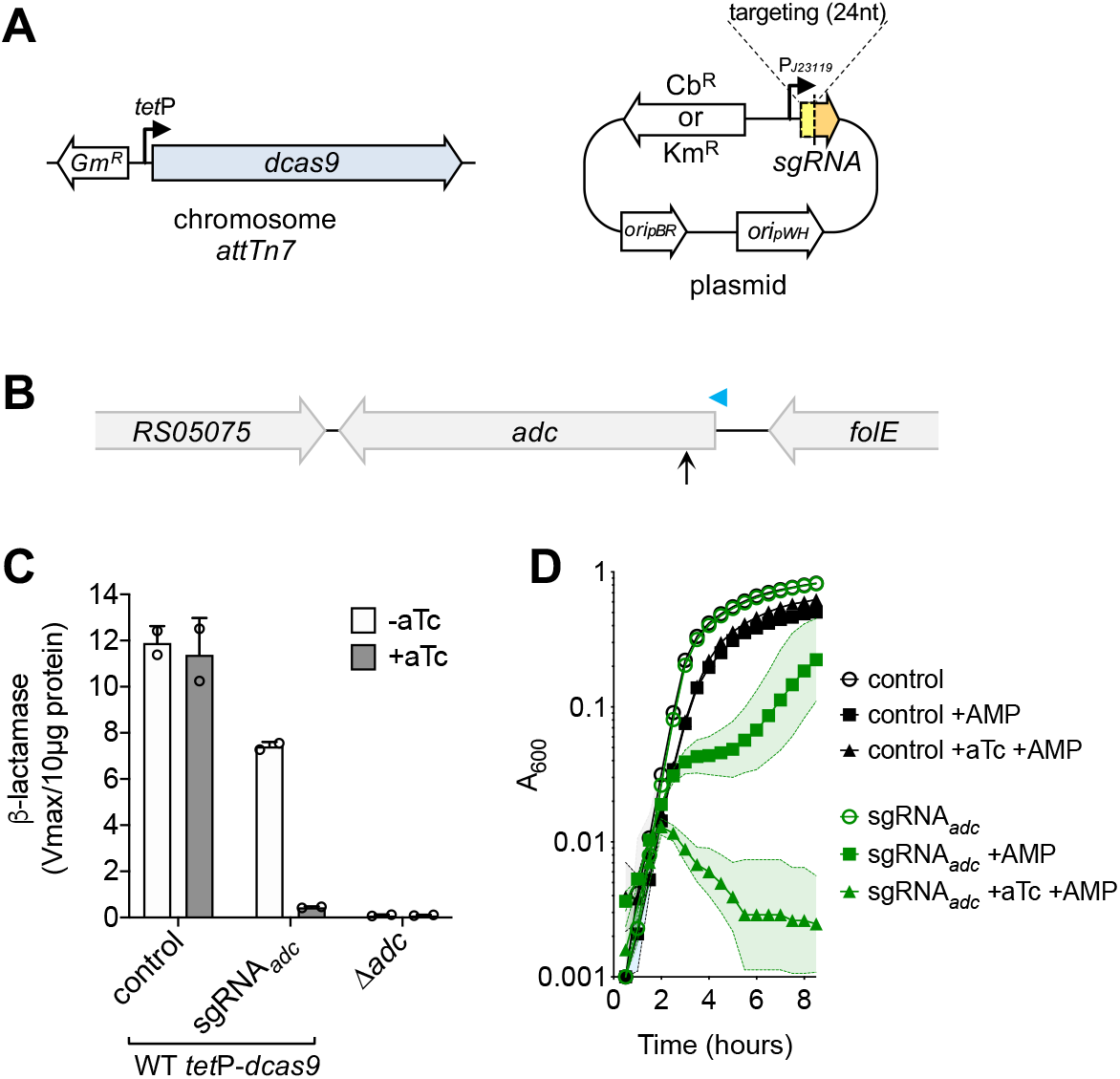
CRISPRi system for efficient knockdown of gene expression in *A. baumannii*. (A) The *A. baumannii* CRISPRi system comprises a chromosomal *dcas9* gene with aTc-inducible expression driven by the *tet* promoter at the *attTn*7 locus, and a high-copy plasmid-based sgRNA transcribed from constitutive J23119 promoter. *ori*pWH indicates pWH1266 origin of replication; *ori*pBR indicates pBR322 origin of replication. (B-D) Efficient CRISPRi-knockdown of the *adc* β-lactamase. (B) Diagram shows a region of the genome containing *adc*. Portions of ACX60_RS05705 and *folE* loci are shown. The position targeted by sgRNA*adc* is indicated by black vertical arrow, and the predicted TSS of *adc* mRNA (23) is shown as blue arrowhead. (C) Cultures of the indicated strain were grown with or without aTc (100 ng/ml) and β-lactamase in sonicates was measured. Bars show mean ± SD (n = 2). (D) *adc* knockdown enhances susceptibility to ampicillin. YDA004 (ATCC 17978 *tet*P-*dcas9*) with sgRNA_*adc*_ or control plasmid was cultured in microtiter format in the presence or absence of aTc (100 ng/ml) and/or ampicillin (AMP, 16 μg/ml) and growth monitored by optical density measurements. Symbols indicate geometric mean and area-filled dotted bands indicate SD (n = 3). Where not visible, SD is within the confines of the symbol. Control refers to pYDE007 (non-targeting control plasmid).

We next used CRISPRi to examine essential genes. We first focused on key division proteins FtsZ and AdvA. The latter is a newly identified protein that plays an essential function in cell division in *A. baumannii*, shown by phenotypic analysis to participate in coordinating chromosome segregation with cell division (5). Viable transposon insertions were almost completely undetectable in *ftsZ*, as predicted for this essential gene, and were detected only within a narrow, central region of *advA*, in agreement with previous findings with lower-density mutant banks (Fig. 3A) (5). Insertions within this region, which may represent a nonessential linker between two essential AdvA domains not tolerating mutation, likely contributed to conflicting essentiality calls (*mariner* predicted essential; Tn*10*-ATS predicted non-essential; Table S2). With CRISPRi constructs targeting *ftsZ* (NT strand, starting 79 bp downstream of predicted TSS(25), Table S2), uninduced cells grew rapidly as short rods characteristic of WT *A. baumannii*, while *dcas9* induction with aTc (200 ng/ml) blocked growth and resulted in a filamentous morphology after 3 hours (Fig. 3B, C). This level of *dcas9* induction had no significant effect on growth or morphology with a non-targeting control sgRNA (Fig. S2). We have shown previously using a conditional allele that *advA* deficiency inhibits growth (5). To confirm this phenotype by CRISPRi, we designed an sgRNA construct that targeted the NT strand of its coding region starting 132 bp downstream of the nearest predicted TSS (Fig. 3A, Table S2). The targeted region was at least 60bp away from the predicted TSS of the divergently transcribed neighboring gene, *serB*, outside of the region bound by the initial RNAP complex (−55 to +20 from a TSS); therefore, this site should not block *serB* transcription (11). As with *ftsZ*, this *advA-*targeting sgRNA resulted in growth inhibition (Fig. 3D) and striking filamentation after 3 hours of *dcas9* induction (Fig. 3E). While growth inhibition occurred less rapidly and at a higher cell density than with *ftsZ*, *advA* knockdown cultures were completely blocked for growth when back-diluted to lower cell density in fresh induction medium (Fig. 3D). These results support our previous genetic analysis demonstrating an essential role for AdvA in *A. baumannii* cell replication (5).

**Fig. 3.**
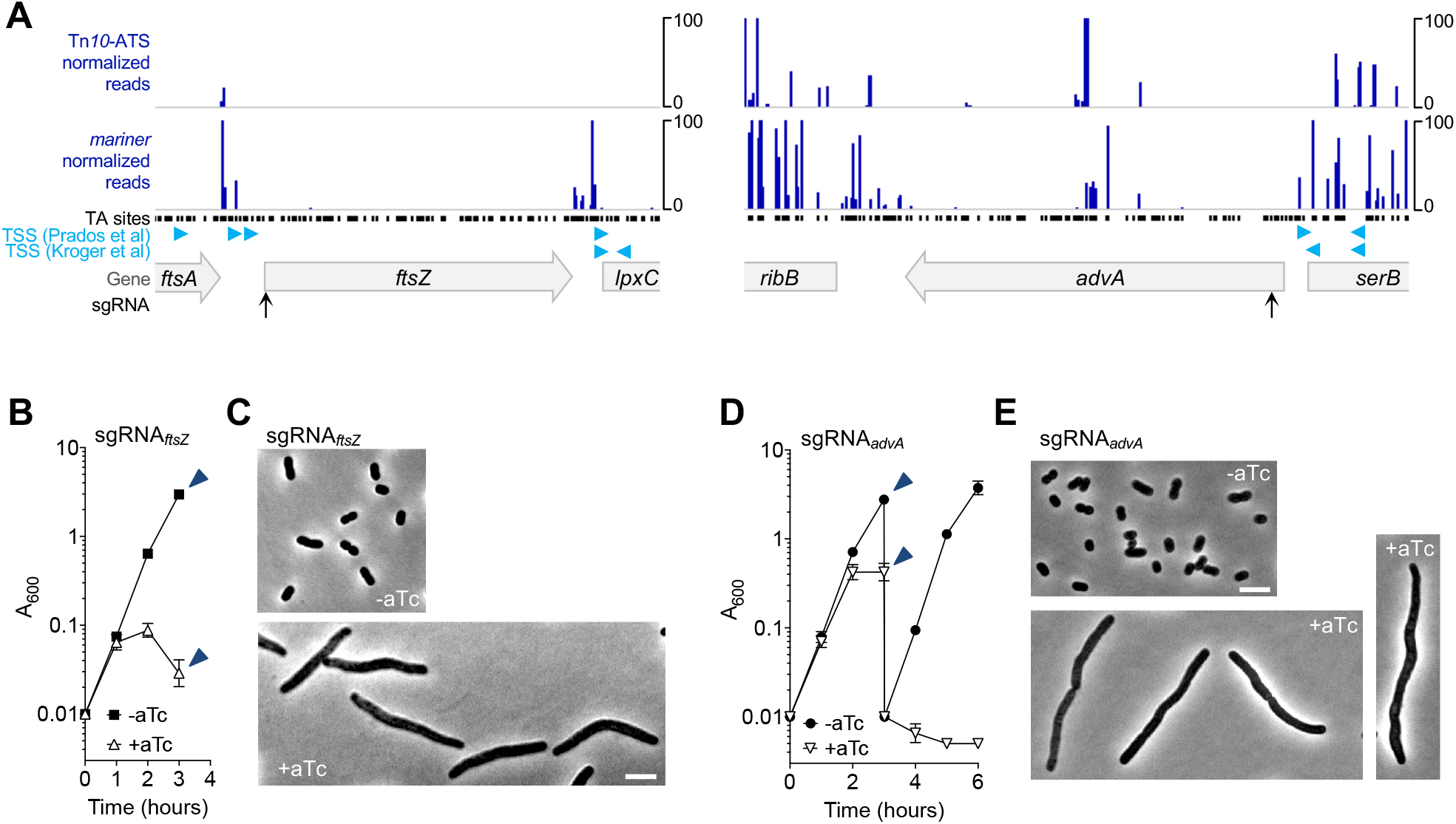
CRISPRi knockdown of *ftsZ* and *advA* blocks growth and cell division. (A) Transposon mutations in *ftsZ* and in most regions of *advA* were not detectable within Tn-seq pools. Normalized Tn-seq read counts were plotted according to position of their transposon insertion in *mariner* and Tn*10*-ATS libraries. Locations of potential *mariner* insertion sites (TA dinucleotides) within these regions are plotted as black points. Predicted TSSs (23, 25) are shown as blue arrowheads. Position targeted by sgRNAs are indicated by vertical black arrows. (B) YDA004 with sgRNA_*ftsZ*_ was cultured with or without 200 ng/ml aTc, and growth was monitored by A_600_ measurements. Data points show geometric mean ± SD (n = 3). At 3 hours, samples were collected (arrowheads) and cells were imaged by phase-contrast microscopy (C). (D, E) AdvA is essential for growth in LB. (D) YDA004 with sgRNA_*advA*_ was cultured for 3 hours in LB with or without 200 ng/ml aTc, then back-diluted to A_600_ = 0.01 into the same (fresh) medium and cultured for an additional 3 hours. Growth was monitored by A_600_ measurements. Data points show geometric mean ± SD (n = 3). Samples were taken at the time points indicated by arrowheads in and cells were imaged with phase contrast (F). Scale bars = 5 μm.

### ACX60_RS03245, encoding a predicted transcription factor, is an essential gene

Transcriptional regulation in *A. baumannii* shows some unusual features, including a small number of sigma factors (4), a large number of transcriptional regulators that jointly control virulence and antibiotic resistance (22, 26, 27), and a reliance on the transcriptional regulator Hfq for growth (28, 29). Given these features and its divergence from other *Gammaproteobacteria*, we predicted that the pathogen may encode unidentified essential transcriptional regulators. As shown in Table S3, we identified 11 candidate essential transcription factors from our Tn-seq datasets. These included the housekeeping sigma factor (*rpoD*), the heat-shock sigma factor (*rpoH*) (30), and *hfq,* as expected; two loci encoding proteins with LexA or Cro/CI homology that were internal to prophages (13) (ACX60_RS07435, ACX60_RS10145) and likely suppress lytic phage replication; and several previously uncharacterized putative transcription factors belonging to the TetR or AraC family (Table S3). In addition, we identified *ompR*, a non-essential gene in strain AB5075 (31), as a candidate essential transcription factor in ATCC 17978. Our Tn-seq data are thus able to define candidate transcription factors which may exert control over essential aspects of *A. baumannii* growth.

We focused our analysis on ACX60_RS03245 (hereafter referred to *RS03245*), a previously uncharacterized candidate essential gene encoding a predicted AraC-family transcription factor. *RS03245* is conserved across *A. baumannii* isolates (32) and is also a candidate essential hit in previous Tn-seq analyses (18, 19). AraC-family regulators typically control catabolism of sugars and amino acids, stress responses, and production of virulence factors (33, 34), and it is unusual for a protein of this family to be essential in rich medium. The domain architecture of the *RS03245*-encoded protein is similar to that of other members of the AraC regulator family (Fig. S3A). In addition, structural homology modeling (35) predicted a relationship with CdpR, a nonessential AraC-family regulator that controls quorum sensing and virulence in *Pseudomonas aeruginosa* (36), despite low sequence identity (22%) (Fig. S3B,C). Transposon insertions were undetectable in the *RS03245* locus (Fig. 4A), consistent with the encoded protein playing an important role in controlling processes essential for *A. baumannii* growth.

**Fig. 4.**
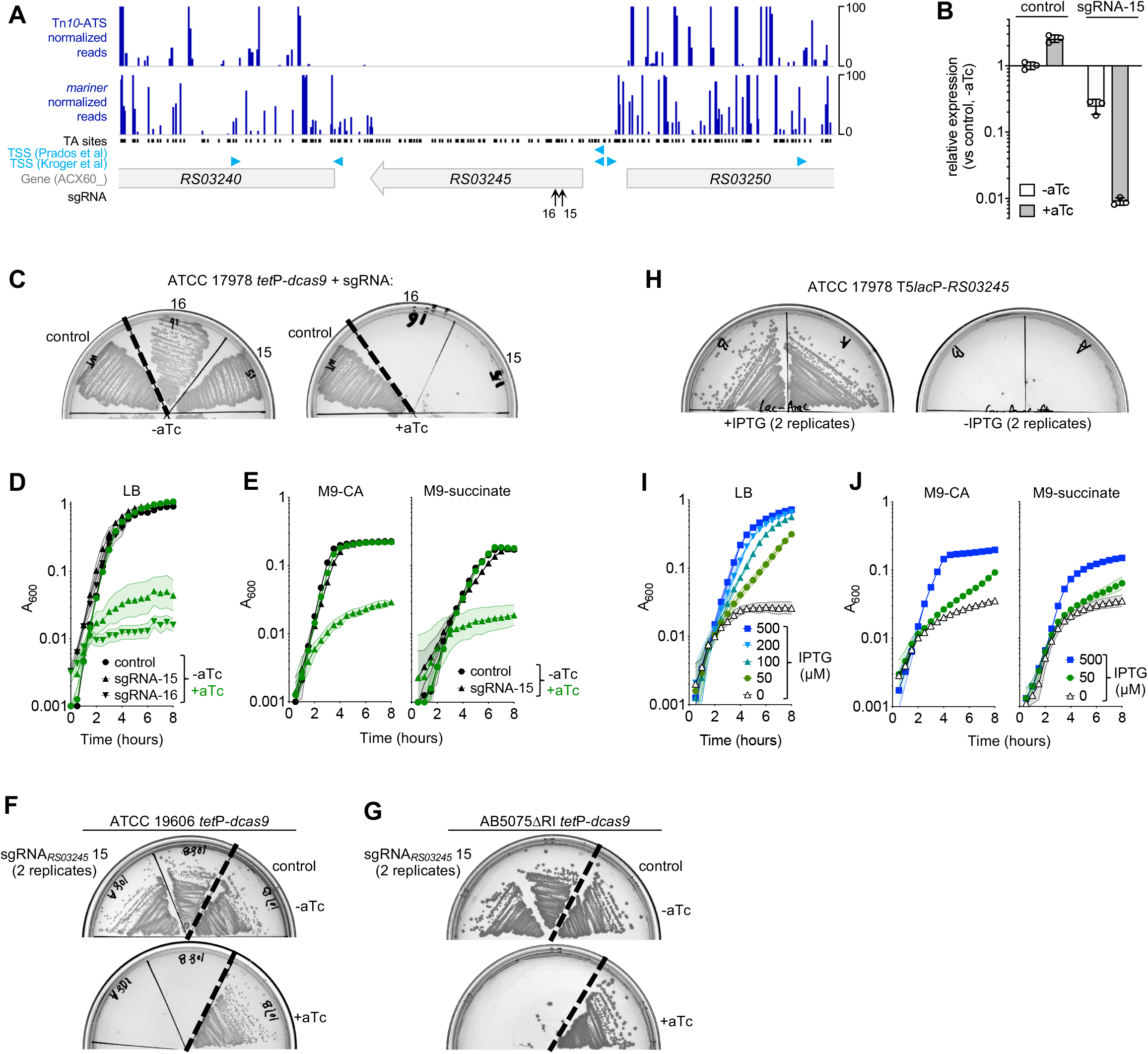
ACX60_RS03245 (*RS03245*) is an essential predicted transcription factor. (A) Transposon mutations within *RS03245* were not detectable in Tn-seq pools. Normalized read counts associated with mutants containing transposons in the region of the *RS03245* locus were plotted according to position of their transposon insertion. Locations of potential *mariner* insertion sites (TA sites), predicted TSSs (23, 25), and sgRNA-targeted positions are indicated as in Fig. 3. (B) Efficient CRISPRi knockdown of *RS03245*. YDA004 with sgRNA targeting *RS03245* (sgRNA-15) or control plasmid was cultured for 2 hours with aTc (50 ng/ml) or no inducer. RNA was extracted and reverse-transcribed, and *RS03245* transcript levels were measured via qRT-PCR. Bars show geometric mean ± SD (n = 3). (C-E) CRISPRi knockdown of *RS03245* blocks growth. YDA004 with indicated sgRNA or control plasmid was grown on solid LB agar ± aTc 200 ng/ml (C) or the indicated liquid medium ± aTc 100 ng/ml in microtiter format (D, E). Data points show geometric mean A_600_ ± SD (n ≥ 2). (F-H) Bacteria containing IPTG-regulated *RS03245* depend on IPTG for growth. JBA58 (T5*lac*p-*RS03245*) was grown on LB agar medium ± IPTG (1 mM) (F) or in microtiter wells with the indicated liquid medium containing IPTG at the indicated concentration (G, H). Data points show geometric mean A_600_ ± SD (n ≥ 3). (I, J) *RS03245* is essential in two additional *A. baumannii* strain backgrounds, ATCC 19606 and AB5075. ATCC 19606 *tet*P-*dcas9* (I) or AB5075ΔRI *tet*P-*dcas9* (J) harboring pJE15 or control plasmid was streaked on solid LB agar without or with aTc (200 ng/μl) and imaged after overnight incubation.

We used CRISPRi to examine the essentiality of *RS03245* predicted by Tn-seq. We designed two separate constructs, sgRNA-15 and sgRNA-16, that target the 5’ end of the *RS03245* coding region near the predicted TSS (Fig. 4A, Table S2). With sgRNA-15, we determined knockdown efficiency by measuring *RS03245* transcription via qRT-PCR. In the absence of *dcas9* induction, *RS03245* transcript levels were decreased 4-fold compared to non-targeting control, while induction for 2 hours with 50 ng/ml aTc caused transcript levels to decrease by more than 100-fold (Fig. 4B). CRISPRi knockdown of *RS03245* is therefore highly efficient. With both sgRNA-15 and sgRNA-16, *dcas9* induction blocked growth on solid (Fig. 4C) and liquid (Fig. 4D) LB medium. Colony formation on LB agar was extremely sensitive to the dose of inducer and was inhibited by sgRNA-15 at aTc concentrations as low as 1.56 ng/ml (Fig. S3D). Growth was also blocked in two types of M9 minimal medium containing either glucose/casamino acids or succinate (Fig. 4E), indicating that *RS03245* essentiality was a general phenotype not specifically depending on rapid growth in rich medium.

We confirmed the above phenotypes by (1) using CRISPRi in two different strain backgrounds and (2) by analyzing *RS03245* essentiality with a completely different genetic approach using a conditional allele. First, we moved the *tet*P-*dcas9* module to the *attTn7* site of *A. baumannii* strains ATCC 19606 and AB5075∆RI, a derivative of AB5075 lacking two large resistance islands (37). *dcas9* induction in the presence of an *RS03245*-targeting guide (sgRNA-15) but not the control construct completely blocked colony formation on LB agar medium with both strain backgrounds (Fig. 4F, G), indicating that *RS03245* has an important function independent of strain background. This is supported by the finding that *RS03245* was an essential gene candidate with AB5075 based on Tn-seq (19). Second, we engineered a derivative of ATCC 17978 in which *RS03245* expression was IPTG-dependent. This was accomplished by replacing the *RS03245* promoter with a *lacI*^q^-T5*lac*P control module using homologous recombination. This mutant (JBA58) depended on IPTG for growth on LB agar (Fig. 4H), in liquid LB (Fig. 4I), as well as in minimal M9 media (Fig. 4J), with growth increasing with increasing IPTG concentration (Fig. 4I,J). These phenotypes closely resembled the effects of CRISPRi knockdown of *RS03245*. Introducing a constitutive copy of *RS03245* (as fusion to either GFP or 3XFLAG epitope) into JBA58 restored the ability to grow in the absence of IPTG (Fig. S3E), indicating that the IPTG-dependence growth phenotypes could be attributed solely to control of *RS03245* expression. Together these results establish *RS03245*, encoding a predicted AraC-family transcription factor, as a novel essential gene in *A. baumannii*.

### Conditional essentiality of late-stage capsule biosynthesis proteins

The consequences of blocking synthesis of capsule, a key virulence factor, on the physiology of *A. baumannii* is incompletely understood (3). Based on bioinformatics analyses and homology with well-studied systems in other organisms (38, 39), capsule biosynthesis across diverse *A. baumannii* isolates is by a Wzy-dependent pathway in which activated sugars (Fig. 5A, steps encoded by genes shaded purple and blue) are utilized by sequential glycosyltransferases (Fig. 5A, encoded by genes shaded green) to build an oligosaccharide repeat unit on an undecaprenyl phosphate (Und-P) lipid carrier (40). The repeat units are then flipped, polymerized and exported to the surface (3) (Fig. 5A, steps encoded by genes in orange). In addition to preventing the formation of structural capsule, defects in this pathway occurring after the initial glycosyltransferase step (Fig. 5A, pink) may have toxic consequences if they generate stalled intermediates that sequester Und-P, which is essential to peptidoglycan synthesis (3, 41, 42). In our high-density Tn-seq analysis, 7 out of 9 genes encoding enzymes predicted to act after the ItrA initiating glycosyltransferase were candidate essential genes (Table S1, *gtr7*, *gtr8*, *wzx*, *wzy*, *wza*, *wzb*, and *wzc*). This analysis is consistent with the model that lesions in late steps in capsule synthesis acting after a committed step are lethal due to the generation of dead-end intermediates.

**Fig. 5.**
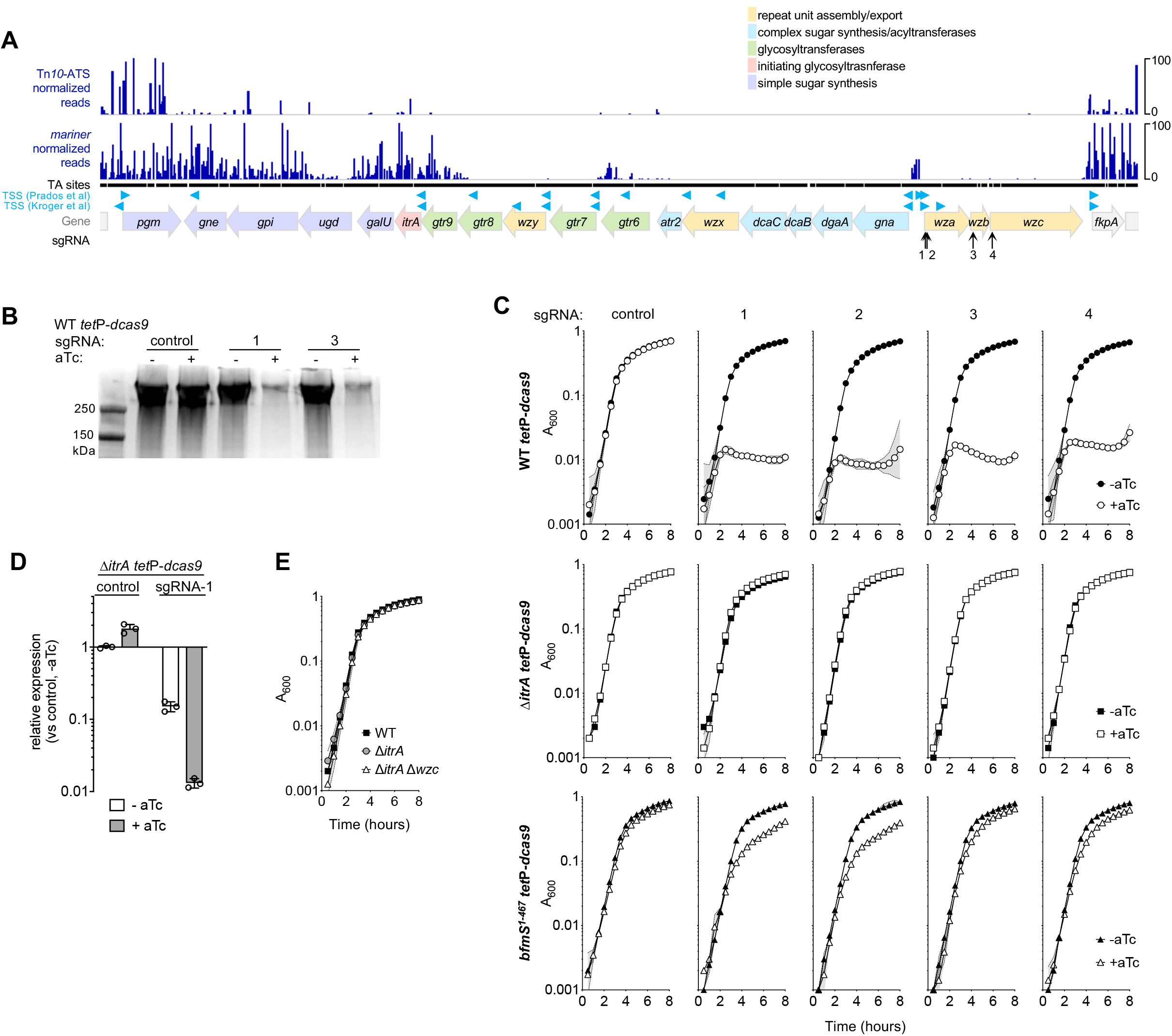
Conditional essentiality of capsule biosynthesis. (A) Location and associated Tn-seq read abundance of transposon mutations within the K locus. Normalized read counts, predicted TSSs (23, 25), and sites targeted by sgRNAs are shown as in Fig. 3. Locations of potential *mariner* insertion sites (TA sites) are plotted as black points below the normalized read counts and appear merged due to the wide view (approximately 25 kb). (B) CRISPRi knockdown of *wza-wzb-wzc* transcriptional unit with the indicated sgRNA inhibits production of capsule. Capsular polysaccharide was analyzed in cell lysates by SDS-PAGE with alcian blue staining. Cultures were grown without or with aTc (200 ng/ml). (C) CRISPRi knockdown of late-stage capsule export blocks growth in a WT strain background (top, circles), but growth inhibition is suppressed with *itrA* deletion (middle, squares) or *bfmS* mutation (bottom, triangles). Strains containing the indicated sgRNA were grown in microtiter format in LB ± aTc 200 ng/ml. Data points show geometric mean A_600_ ± SD (n ≥ 2). Δ*itrA tet*P-*dcas9* strain was JBA9; *bfmS*^1−467^ *tet*P-*dcas9* strain was JBA8. (D) CRISPRi knockdown with sgRNA-1 (targeting 5’ end of *wza*) blocks *wza* transcription in Δ*itrA* background. Strains were grown with the indicated inducer concentration for 1 hr. RNA was extracted, reverse-transcribed and *wza* transcript levels probed via qPCR. Shown is the mean of the fold change in transcript levels vs control, untreated ± s.d. (n = 3). P < 0.04 in unpaired t-tests comparing sgRNA-1 vs control with each treatment condition. (E) Δ*wza* Δ*itrA* double mutant was easily isolated (JBA48) and shows growth kinetics identical to that of its Δ*itrA* parent (EGA295) and WT. Data points show geometric mean A_600_ ± SD (n = 3).

We employed CRISPRi to test this model and assess whether block of the early step, by ItrA, in which Und-P acceptors are likely dedicated to capsule can relieve toxicity. First, we confirmed that knockdown of the late-stage capsule export module (encoded by the co-transcribed genes *wza*-*wzb*-*wzc*, Fig. 5A) prevents growth. We designed 4 distinct sgRNA constructs targeting different positions within this operon (Fig. 5A, vertical arrows; Table S2). While CRISPRi has polar effects and these sgRNAs likely modulate the entire *wza*-*wzb*-*wzc* operon, each gene encodes a part of the same complex dedicated to capsule export and high-level polymerization (43). Interference of operon transcription by CRISPRi should thus enable targeted examination of this process. With two of the sgRNAs (1 and 3), we verified via alcian blue staining of SDS-PAGE-separated cell lysates ((44) and Materials and Methods) that production of capsular polysaccharides was blocked after *dcas9* induction (Fig. 5B). In the absence of inducer these sgRNAs had minimal effect on capsule production (Fig. 5B). Consistent with these results and our Tn-seq analysis, sgRNA-1 through 4 each blocked growth only when *dcas9* was induced (Fig. 5C, circles). To test the model that late-step capsule synthesis defects can be tolerated when the predicted committed step in Und-P usage is prevented (by blocking ItrA), we moved the CRISPRi machinery into EGA295 (ATCC 17978 ∆*itrA*) and repeated the *dcas9* induction experiment. Strikingly, CRISPRi block of *wza*-*wzb*-*wzc* had no effect on growth in the absence of *itrA* (Fig. 5C, squares), despite knockdown being highly efficient under these conditions as determined by measuring *wza* transcription levels (Fig. 5D). The ability of Δ*itrA* to suppress the nonviability caused by late-stage capsule block was confirmed by isolating a Δ*itrA* Δ*wzc* double mutant strain. Unlike the Δ*wzc* single mutant which could not be successfully isolated from an ItrA^+^ background in the absence of compensatory mutations (44), Δ*wzc* was easily isolated in a Δ*itrA* background, and the resulting double mutant had WT growth kinetics (Fig. 5E). Together these results indicate that the essentiality of late steps in capsule assembly can be bypassed when flux into the pathway is prevented.

We used CRISPRi to confirm an additional pathway allowing bypass of lethality associated with capsule production defects. Our previous work identified *bfmS* as a site of suppressor mutations that allowed growth of a Δ*wzc* strain (44). BfmS is part of the BfmRS two-component system, and null mutations in *bfmS* cause augmented expression of envelope stress response genes and genes determining synthesis of envelope structures including capsule and Und-P (22). These changes may enhance how cells cope with deleterious dead-end intermediates associated with capsule production defects. We used CRISPRi to confirm the suppressive interaction between *bfmS* and late-stage capsule block. After moving the CRISPRi system to a BfmS^−^ strain background, (*bfmS*^1−467^, Table S2, (22, 44)), guides 1-4 were tested for ability to inhibit growth in the absence and presence of *dcas9* induction. As predicted and in contrast to WT (BfmS^+^) bacteria, *bfmS*^1-467^ cells tolerated CRISPRi targeting of *wza*-*wzb*-wzc despite *dcas9* induction, although partial growth inhibition was observed compared to the control sgRNA (Fig. 5C, triangles). These results are consistent with envelope stress response activation providing a second pathway to allow toleration of late stage capsule synthesis defects.

## Discussion

In this study, we have defined a candidate essential gene set in *A. baumannii* and we have established a CRISPRi tool for analysis of these genes. Our essential gene search was comprehensive and utilized two different global transposition systems based on unrelated classes of transposase enzymes. This approach provided a high level of saturation of the genome and allowed us to build a consensus set of candidate essential genes. This candidate essential gene set can guide the selection of targets in future studies aimed at dissecting the *A. baumannii* essential genome.

To target essential genes, we developed a CRISPRi knockdown system for *A. baumannii*. The system was controllable and efficient, with low-level (~2-4 fold) decreases observed in the absence of *dcas9* induction by aTc, and effective shut-down of gene expression (by ~30- to 100-fold with different genes) with induction. Knockdown is likely to be titratable using dilutions of aTc, as indicated by the relationship of aTc concentration with colony formation by cells containing *RS03245*-targeting sgRNA (Fig. S3D). CRISPRi knockdown was as effective as classical allelic replacement techniques in allowing analysis of conditional growth phenotypes and was functional in multiple strain backgrounds (Fig. 4). The efficiency of our system is an advancement over a previous system which showed ~10-fold knockdown with *A. baumannii* (12). By using pools of diverse guides, this system should facilitate large-scale examination of terminal and hypomorphic phenotypes linked to essential genes (11, 45). Such studies have the potential to illuminate essential protein function and inform novel antimicrobial target development, and would complement previous functional genomics studies that focused largely on nonessential genes in *A. baumannii*(5).

The candidate essential gene set and knockdown system developed in this study facilitated the confirmation that *advA* is an essential gene in *A. baumannii*, identification of essential predicted transcription factors including *RS03245*, and demonstration of the conditional essentiality of capsule export proteins. Part of the core genome (32), *RS03245* is one of 33 genes encoding AraC-family transcription factors in *A. baumannii* but was the only one determined to be essential in this and previous Tn-seq studies (Table S4). By analogy with most AraC-family proteins which are transcriptional activators (33), *RS03245* may enhance transcription of genes contributing to one or more essential pathways. Studies are underway to characterize the *RS03245* regulon and its relationship to essential genetic networks in the organism. In addition, we demonstrated that late steps in capsule biosynthesis are essential in ATCC 17978 unless suppressed by one of at least two pathways controlled by *itrA* or *bfmS*. This conditional essentiality mirrors that seen with defects in Wzy-dependent synthesis of capsule in *S. pneumoniae* (46, 47) and *E. coli* (48), and of O-antigen in *E. coli* (49). It also parallels the conditional toxicity of LOS synthesis lesions in *A. baumannii*, in which defects in late-acting synthesis steps are lethal unless flux into the pathway is reduced (50–52). The molecular basis for toxicity and for the differential ability of *A. baumannii* strains to cope with capsule synthesis defects (3) is unclear. Understanding these processes in future studies may inform strategies to attack the pathogen’s protective envelope.

In summary, we have identified candidate essential genes in *A. baumannii* and developed a CRISPRi-based tool that facilitated rapid validation of the essentiality of a number of these candidates. This tool should enhance both targeted and large-scale analyses of essential processes in the microorganism. In addition, the essential pathways determined in this work open new avenues for research into the pathogen’s dependence on transcriptional control and envelope homeostasis for growth and survival.

## Materials and Methods

### Bacterial strains, growth conditions, and antibiotics

Bacterial strains used in this work are described in Table S5. *A. baumannii* strains were derivatives of ATCC 17978 unless otherwise noted. Bacteria were cultured in Lysogeny Broth (LB) (10 g/L tryptone, 5 g/L yeast extract, 10 g/L NaCl) unless otherwise noted. Cultures were incubated at 37°C in flasks with shaking or in tubes on a roller drum. Growth was monitored by measuring absorbance at 600nm via a spectrophotometer. Where indicated, microtiter format growth was with 96-well plates incubated with shaking in a plate reader (Epoch 2 or Synergy H2M, Biotek). LB agar was supplemented with antibiotics [carbenicillin (Cb) at 50-100 μg/ml with all strains except *A. baumannii* AB5075ΔRI, 1600μg/ml), kanamycin (Km) at 10-20 μg/ml, gentamicin (Gm) at 10 or 40μg/ml] or sucrose as needed (Sigma Aldrich).

### Construction of transposon mutant libraries

Mutagenesis with the *mariner* and Tn*10*-ATS transposon systems was performed by electroporation with pDL1100 and pDL1073, respectively (5, 13). Transformed cells were spread on membrane filters, allowed to recover on solid SOC, and enriched after transfer to selective solid LB-Km medium as described (5, 13). Colonies were lifted from filters by agitation in sterile PBS, mixed with sterile glycerol (10%), aliquoted, and stored at −80°C. With *mariner*, colonies from 10 previously constructed *mariner* subpools (5) as well as 28 additional subpools constructed for this study were analyzed in aggregate, representing a total of approximately 550,000 mutant colonies. With Tn*10*-ATS, colonies from 11 previously constructed subpools (13), as well as 12 additional subpools constructed for this study were analyzed in aggregate, representing approximately 300,000 mutant colonies in total.

### Tn-seq Illumina library preparation and sequencing

Genomic DNA was extracted from samples (Qiagen DNeasy Kit) and quantified by a SYBR green microtiter assay. Transposon-adjacent DNA was tagmented and amplified for Illumina sequencing using a modified Nextera™ DNA Library Prep method as described (13). Samples were multiplexed, reconditioned, and size selected (250- or 275-600bp, Pippin HT) before sequencing (single-end 50bp) using custom primers (5, 13) on a HiSeq2500 with High Output V4 chemistry at Tufts University Genomics Core Facility.

### Tn-seq data analysis

Sequencing reads were quality-filtered and clipped of adapters with BBDuk, collapsed with fastx_collapser, and mapped to the *A. baumannii* chromosome (NZ_CP012004) with Bowtie (5). To process the *mariner* dataset into an input file for TRANSIT analysis, the coordinates of TA sites in the NZ_CP012004 genome that can be uniquely mapped to (i.e., not part of repeat regions) were identified, and mapped reads were tabulated according to these sites in wig format using custom python scripts. With the Tn*10*-ATS data set, mapped reads were tabulated in a wig file containing all chromosome coordinates using python. Read counts were normalized across all subpools within a dataset (*mariner* or Tn*10*-ATS) by the TTR method with the TRANSIT software package (16) and merged into a single wig file for each dataset, and scaled such that median read coverage at non-zero insertion sites was similar between datasets. To determine gene essentiality with TRANSIT, the *mariner* dataset was analyzed by the Gumbel method (parameters: ignore C-terminal 10%, Sample Size 10000, Burn-in 500, Trim 1, minimum read 5), and Tn*10*-ATS was analyzed by Tn5gaps method (parameters: ignore C-terminal 10%, minimum read 5) (16). Orthologs in CP000521 (ATCC 17978 genome file used in (18)) and CP008706 (AB5075-UW genome used in (19)) were matched to NZ_CP012004 genes by using Mauve to identify positional orthologs (53) and by Boundary-Forest Clustering (54). Relationship of essential genes were analyzed via Venn diagrams using BioVenn (55). Integrative Genomics Viewer (56) was used to visualize normalized Tn-seq read counts and unique TA insertion sites along chromosome coordinate.

### Molecular cloning and mutant construction

Oligonucleotide primers and plasmids used in this study are listed in Table S6. All constructs containing cloned PCR products were verified by sequencing (Genewiz). A miniTn*7* element containing *dcas9* was constructed by PCR-amplifying the *tetR*-*tet*P-*dcas9*-*rrnB*T1-T7Te fragment from pdCas9-bacteria (Addgene #44249, gift of Stanley Qi) with primers containing SpeI and PstI sites, cloning in the HincII site of pUC18, and subcloning in the SpeI and PstI sites of pUC18T-miniTn7T-Gm (57) to generate pYDE009 (Cb^r^, Gm^r^). The miniTn*7* element of pYDE009 was moved into *A. baumannii* by four-parental mating (44, 58) using Vogel Bonner Medium with Gm at 10 μg/ml (ATCC 17978, AB5075ΔRI) or 40μg/ml (ATCC 19606), creating YDA004 and JBA106, respectively.

Integration at the *attTn7* locus was confirmed by PCR (20, 59). A plasmid for sgRNA expression in *A. baumannii* was constructed by replacing the BamHI-SalI fragment internal to *tetA* in shuttle vector pWH1266(21) with a PCR product having the same restriction sites and J23119, sgRNA, and terminator from pgRNA-bacteria (Addgene #44251, gift of Stanley Qi), generating pYDE007 (Cb^r^). To construct a Km^r^ derivative of pYDE007, the EcoRI-PstI fragment containing *bla* was replaced with a PCR product containing the kanamycin resistance gene from pDL1100 and the same restriction sites, generating pJE53 (Km^r^). pYDE0007 and pJE53 contain the non-targeting guide sequence, AACTTTCAGTTTAGCGGTCT, derived from the mRFP guide in pgRNA-bacteria. These plasmids served as non-targeting controls in CRISPRi experiments.

Plasmids encoding sgRNAs for targeted CRISPRi were constructed by PCR-amplifying the gRNA scaffold region of pYDE007, using a forward primer containing a 24-base targeting sequence and SpeI site at its 5’ end, a reverse primer with ApaI in its 5’ end, and OneTaq PCR Master Mix (NEB). Reverse primers for most sgRNAs also contained a unique KpnI or BglII site to assist clone identification by restriction fragment analysis. PCR products were digested with SpeI and ApaI and cloned by replacing the SpeI-ApaI guide fragment in pYDE007. sgRNA plasmids were introduced into YDA004 and JBA106 via electroporation. sgRNA targeting sequences were the 23 or 24 bases 5’ to PAM sites selected based on (i) proximity to the TSS preceding each target gene, (ii) targeting the non-template strand, and (iii) having a 12-nt seed region found only once in the genome (11, 60).

A strain containing a conditional allele of *RS03245* was constructed as follows. *RS03245* was first cloned as a translational fusion to GFP in pJC180 using BamHI and XbaI sites, and the fusion gene was subcloned in pYDE152 using SmaI and PstI sites, generating *lacI*^q^-T5*lac*P-*RS03245*-*gfp* (pJE10). As 3’ homology arm, the *lacI*^q^-T5*lac*P-*RS03245* sequence was PCR amplified with primers containing SphI and NotI sites and cloned in pUC18, generating pJE41. As 5’ homology arm, approximately 1kb of sequence upstream of *RS03245* was PCR-amplified with primers having SphI and SalI sites and cloned in pUC18, generating pJE40. Clones were joined and subcloned in pJB4648 by 3-way ligation, generating pJE44. pJE44 was delivered into *A. baumannii* ATCC 17978 by electroporation, and counterselection allelic exchange (44) was used to isolate JBA58, in which the native *RS03245* promoter is replaced with *lacI*^q^-T5*lac*P.

To construct complementing plasmids, *RS03245*-GFP or *RS03245* translationally fused to a 3X-FLAG epitope were subcloned between the BamHI and SphI sites internal to *tetA* in pWH1266, generating pJE51 and pJE50, respectively. In each, *RS03245* is expressed by a constitutive *tet* promoter.

A Δ*itrA* Δ*wzc* mutant was constructed by introducing pEGE76 (allelic exchange vector with Δ*wzc*::Gm^r^ (44)) into EGA295 (Δ*itrA* single mutant) by electroporation, and using counterselection allelic exchange to isolate JBA48(Δ*itrA* Δ*wzc*, Gm^r^).

### β-lactamase assays

Overnight bacterial cultures were back-diluted to A_600_ 0.025, grown to A_600_ 0.05, and cultures were divided with one group receiving aTc (100 ng/ml). After 2 additional hours of growth, cells were harvested by centrifugation, washed, and resuspended in ice-cold 0.1 M phosphate buffer (pH 7). Periplasmic contents were liberated by ultrasonication using a Branson high-intensity cuphorn sonifier chilled to 4°C (4 cycles of 1 min ON at 50% output/1 min OFF). Extracts were clarified by centrifugation, diluted with 0.1M phosphate buffer (pH 7), and protein concentration was determined via Bradford assay (Pierce). β-lactamase content was measured in 250 μl reaction mixtures containing 10 μg total protein and 20μg/ml nitrocefin in 0.1M phosphate buffer, pH 7 by reading absorbance at 486 nm every minute for 15 minutes at room temperature in microtiter plates (Biotek Synergy H1M). β-lactamase activity was calculated as initial reaction rate (Vmax) × dilution factor.

### Microscopy

Bacteria were immobilized on agarose pads (1% in PBS), and imaged via 100x/1.3 phase-contrast objective on a Leica AF6000 microscope.

### Analysis of capsular polysaccharide

Overnight bacterial cultures were diluted to OD 0.025, grown to OD 0.05, and divided into groups receiving 0 or 200ng/ml aTc. Cultures were grown for 4 additional hours. Polysaccharides were isolated from whole-cell lysates with slight variations from previously described methods (44, 61). Cells were pelleted, frozen at −80°C, and resuspended with 60mM Tris, pH 8 buffer containing 10mM MgCl_2_ and 50μM CaCl_2_. 0.5 μl lysonase (Novagen) was added per 100 μl suspension, and samples were incubated at 37°C for 30 minutes followed by vortexing. SDS was added to 0.5%, and samples were incubated at 37°C for 15 minutes. 2 μl Proteinase K (NEB) was added and samples were incubated at 56°C for 1 hour. SDS sample buffer was added to 1X concentration and samples were boiled for 5 minutes. Samples were separated on 4-20% BioRad TGX Tris-glycine gels and stained overnight with alcian blue. Gels were imaged via white light transillumination with a ChemiDoc MP (BioRad).

### qPCR gene expression analysis

Overnight bacterial cultures were diluted to OD 0.025, grown to OD 0.05, and divided into aTc-treated or untreated groups. After 2 (sgRNA_*RS03245*_-15) or 4 hours (sgRNA-1) additional growth, samples were combined with one volume of ice-cold ethanol-acetone and frozen at −80°C. Samples were thawed and washed with TE, followed by RNA extraction (RNeasy kit, Qiagen) and DNase treatment (DNA-free kit, Ambion). RNA was reverse transcribed using Superscript II Reverse Transcriptase (Invitrogen). cDNA was diluted and used as template with the Power-Up SYBR Green Master Mix (Applied Biosystems) in a StepOnePlus system according to the manufacturer’s instructions for two-step RT-PCR. Primers targeting *RS03245* and *wza* were designed using PrimerQuest (IDT). Assay efficiency was assessed by generating a standard curve with a dilution series of cDNA and was determined to be >99% with each target. Controls lacking reverse-transcriptase were performed to confirm lack of signal from residual genomic DNA. Gene expression levels were quantified by using the 2^-ΔΔCt^ method with *rpoC* as endogenous control (22).

## Acknowledgements

This work was supported by Northeastern University College of Science startup funds and NIAID awards U01AI124302 and R21AI128328. We thank Stanley Qi for plasmid gifts, Colin Manoil for strain AB5075ΔRI, Elizabeth Schwartz for technical assistance, and members of the Geisinger lab for helpful discussions.

## Supplemental Figure Legends

**Fig. S1.**
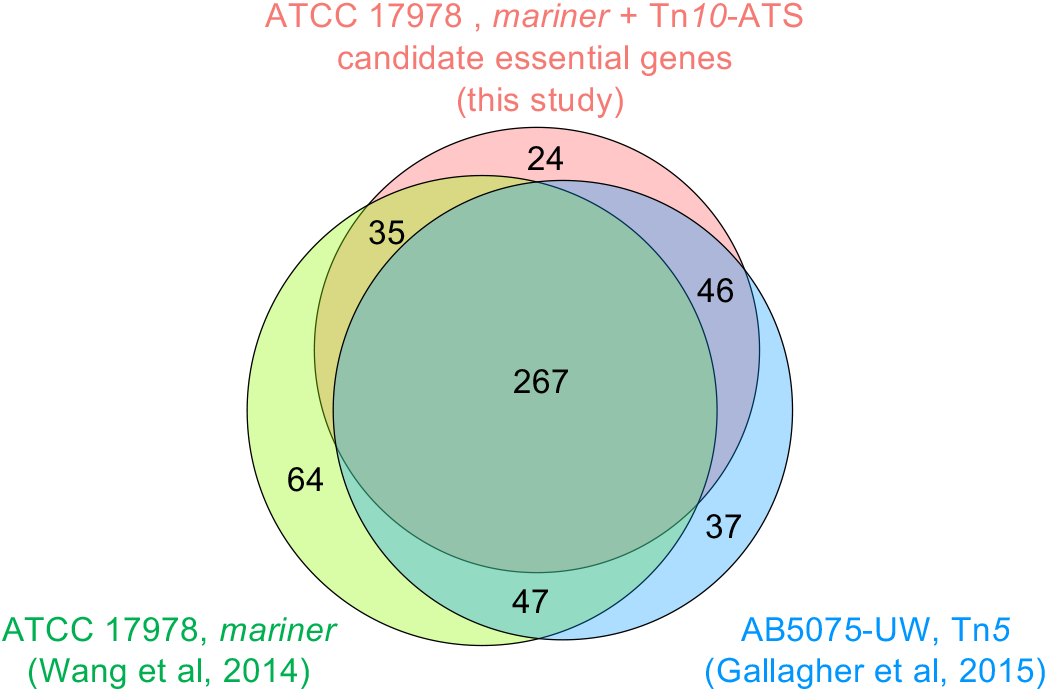
Candidate *A. baumannii* essential genes correspond well with candidates identified in prior studies. Venn diagram shows the relationship of candidate essential genes determined with ATCC 17978 in this study by both *mariner* and Tn*10*-ATS systems (red circle) with sets of candidate essential genes previously identified by Tn-seq in the same strain (18) (green circle) and in strain AB5075-UW (19) (blue circle). Class of transposon utilized in each Tn-seq analysis is indicated.

**Fig. S2.**
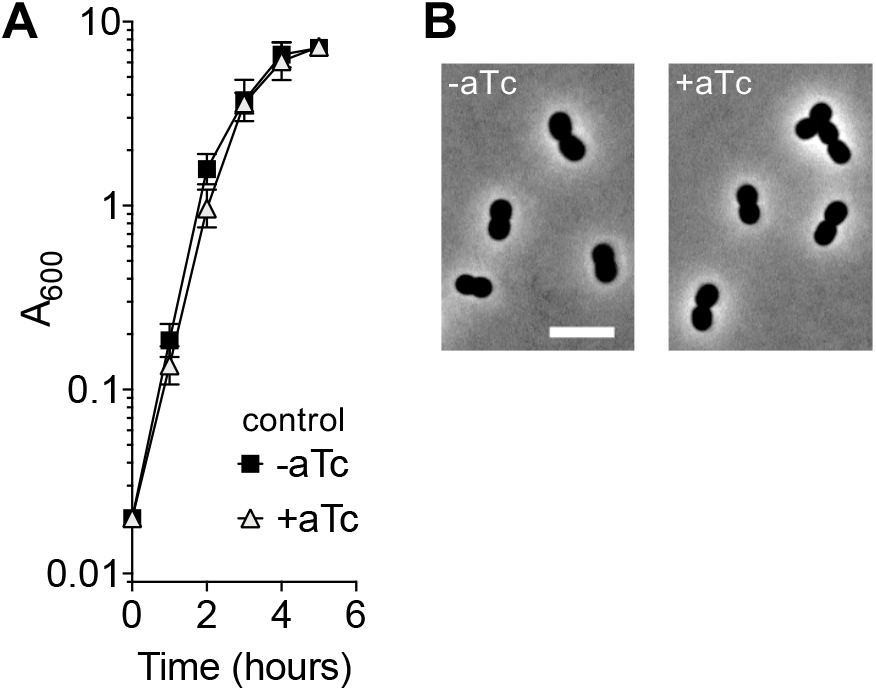
Induction of *dcas9* with control non-targeting sgRNA does not alter growth or morphology. (A) YDA007 (*tet*P-*dcas9*, control sgRNA) was cultured with or without 200 ng/ml aTc, and growth was monitored by A_600_ measurements. Data points show geometric mean ± SD (n = 3). No significant difference was detected at any time point (p > 0.05, t tests). (B) YDA007 cells grown as in A were imaged by phase-contrast microscopy. Scale bar, 5μm.

**Fig. S3.**
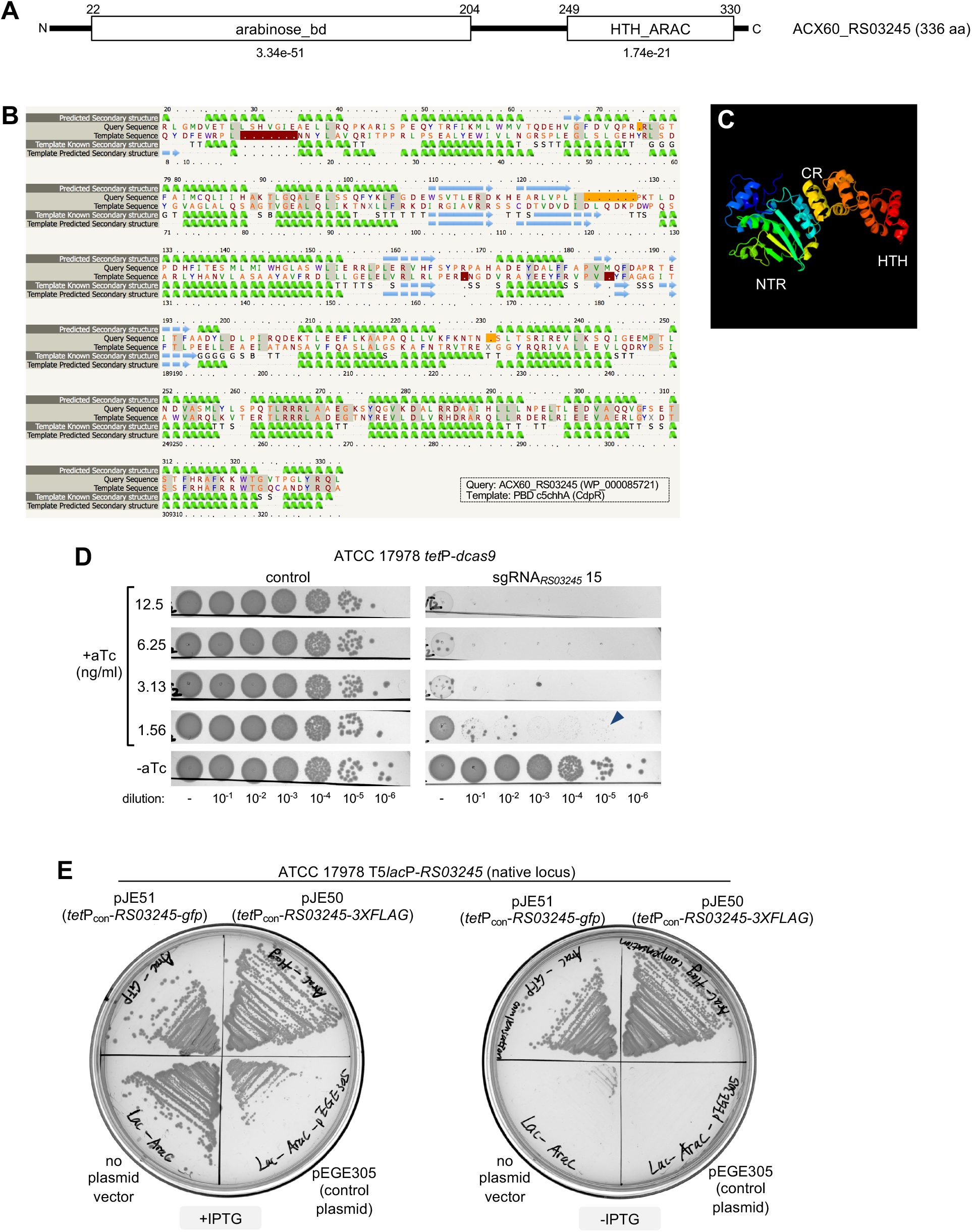
Newly identified essential protein RS03245 shows structural homology with a *P. aeruginosa* transcription factor and essentiality in two *A. baumannii* strain backgrounds. (A) Conserved domains in RS03245 protein (WP_000085721) identified via CDD (62). Numbers above the protein indicate aa residue; numbers below each domain indicate E-value of CDD hit. (B) Sequence alignment between RS03245 protein sequence and known structure of *P. aeruginosa* CdpR (36) via Phyre2 (35). Predicted and known secondary structures are shown. Residues are color-coded based on property as described (35), and identical residues have gray background shading. Confidence of Phyre2 structural homology prediction was 100%, coverage was 90%. (C) 3D Phyre2 model predicting RS03245 protein folding based on structural homology with CdpR. NTR, CR, and HTH domains are indicated based on CdpR (36). (D) *A. baumannii* is highly sensitive to CRISPRi knockdown of *RS03245*. ATCC 17978 *tet*P-*dcas9* harboring pJE15 (sgRNA_*RS03245*_ 15) or control plasmid were cultured in absence of inducer, serially diluted in PBS, and spotted onto solid LB agar containing the indicated aTc inducer concentration. Colonies were imaged after overnight growth. With sgRNA_*RS03245*_ at high cell dilution, pinpoint colonies (arrowhead indicates example) were visible with 1.56 ng/μl, but no colonies were detected at higher inducer concentrations. (E) Rescue of IPTG dependence of T5*lac*P-*RS03245* by cloned *RS03245* expressed from constitutive promoter. ATCC 17978 T5*lac*P-*RS03245* harboring the indicated plasmids were streaked on solid LB agar containing 0 or 1mM IPTG, and imaged after overnight incubation.

**Table S1.** *A. baumannii* gene essentiality determined by Tn-seq.

**Table S2.** sgRNA targeting sequences used in this study.

**Table S3.** Candidate essential transcriptional regulators in *A. baumannii* ATCC 17978

**Table S4.** Essentiality analysis of predicted AraC-family transcription factors encoded in *A. baumannii* ATCC 17978 genome.

**Table S5.** Bacterial strains and plasmids used in this study.

**Table S6.** Oligonucleotide primers used in this study.

