## Supplemental Table 5 for "Essential gene analysis in *Acinetobacter baumannii* by high-density transposon mutagenesis and CRISPR interference"

**Table S5. Strains and plasmids.**

| Strain or plasmid | Genotype or description | Reference |
|---|---|---|
| ***A. baumannii*** | | |
| ATCC 17978 | cerebrospinal fluid isolate | (1) |
| ATCC 19606 | urine isolate | (1) |
| AB5075-UW | bone isolate/osteomyelitis | (2) |
| AB5075ΔRI | AB5075-UW with deletions of the two resistance islands | (3) |
| EGA513 | ATCC 17978 Δ*adc* | (4) |
| EGA295 | ATCC 17978 Δ*itrA* | (5) |
| JBA48 | ATCC 17978 Δ*itrA* Δ*wzc* | This study |
| EGA127 | ATCC 17978 *bfmS*$^{1-467}$(G467DfsX19) | (5) |
| YDA004 | ATCC 17978 *attTn7*::*tet*R-*tet*P-*dcas9*-*rrnB*T1-T7Te Gm$^r$ | This study |
| JBA8 | ATCC 17978 *bfmS*$^{1-467}$ *attTn7*::*tet*R-*tet*P-*dcas9*-*rrnB*T1-T7Te Gm$^r$ | This study |
| JBA9 | ATCC 17978 Δ*itrA* *attTn7*::*tet*R-*tet*P-*dcas9*-*rrnB*T1-T7Te Gm$^r$ | This study |
| JBA106 | ATCC 19606 *attTn7*::*tet*R-*tet*P-*dcas9*-*rrnB*T1-T7Te Gm$^r$ | This study |
| JBA163 | AB5075ΔRI *attTn7*::*tet*R-*tet*P-*dcas9*-*rrnB*T1-T7Te Gm$^r$ | This study |
| YDA007 | YDA004 with pYDE007 (non-targeting control) | This study |
| JBA65 | YDA004 with pJE55 (sgRNA$_{adc}$) | This study |
| YDA009 | YDA004 with pAFE119 (sgRNA$_{ftsZ}$) | This study |
| YDA011 | YDA004 with pYDE317 (sgRNA$_{advA}$) | This study |
| JBA15 | YDA004 with pJE15 (sgRNA$_{\mathrm{RS03245}}$-15) | This study |
| JBA16 | YDA004 with pJE16 (sgRNA$_{\mathrm{RS03245}}$-16) | This study |
| JBA107 | JBA106 with pYDE007 (non-targeting control) | This study |
| JBA108 | JBA106 with pJE15 (sgRNA$_{\mathrm{RS03245}}$-15) | This study |
| JBA166 | JBA163 with pYDE007 (non-targeting control) | This study |
| JBA164 | JBA163 with pJE15 (sgRNA$_{\mathrm{RS03245}}$-15) | This study |
| JBA23 | YDA004 with pJE23 (sgRNA$_{wza\text{-}wzb\text{-}wzc}$-1) | This study |
| JBA22 | YDA004 with pJE22 (sgRNA$_{wza\text{-}wzb\text{-}wzc}$-2) | This study |
| JBA25 | YDA004 with pJE25 (sgRNA$_{wza\text{-}wzb\text{-}wzc}$-3) | This study |
| YDA47 | YDA004 with pYDE318 (sgRNA$_{wza\text{-}wzb\text{-}wzc}$-4) | This study |
| JBA43 | JBA9 with pYDE007 (non-targeting control) | This study |
| JBA41 | JBA9 with pJE23 (sgRNA$_{wza\text{-}wzb\text{-}wzc}$-1) | This study |
| JBA38 | JBA9 with pJE22 (sgRNA$_{wza\text{-}wzb\text{-}wzc}$-2) | This study |
| JBA39 | JBA9 with pJE25 (sgRNA$_{wza\text{-}wzb\text{-}wzc}$-3) | This study |
| JBA40 | JBA9 with pYDE318 (sgRNA$_{wza\text{-}wzb\text{-}wzc}$-4) | This study |
| JBA42 | JBA8 with pYDE007 (non-targeting control) | This study |
| JBA35 | JBA8 with pJE23 (sgRNA$_{wza\text{-}wzb\text{-}wzc}$-1) | This study |
| JBA34 | JBA8 with pJE22 (sgRNA$_{wza\text{-}wzb\text{-}wzc}$-2) | This study |
| JBA36 | JBA8 with pJE25 (sgRNA$_{wza\text{-}wzb\text{-}wzc}$-3) | This study |
| JBA33 | JBA8 with pYDE318 (sgRNA$_{wza\text{-}wzb\text{-}wzc}$-4) | This study |
| JBA58 | ATCC 17978 *lacI*$^q$-T5*lac*P-ACX60_RS03245 | This study |
| JBA64 | JBA58 with pJE51 | This study |
| JBA63 | JBA58 with pJE50 | This study |
| JBA59 | JBA58 with pEGE305 | This study |

| | | |
|---|---|---|
| ***E. coli*** | | |
| DH5α | *supE44* Δ*lacU169* (ϕ80lacZΔM15) *hsdR17 recA1 endA1 gyrA96 thi*-1 *relA1* | (6) |
| DH5λpir | DH5α (λpir) *tet*::Mu *recA* | (7) |
| XL1-blue | *recA1 endA1 gyrA96 thi-1 hsdR17 supE44 relA1 lac* [F′ *proAB lacI*$^{q}$ZΔM15 Tn10 Tc$^{r}$)] | Stratagene |
| **plasmids** | | |
| pDL1100 | *Himar1 mariner* (Km$^{r}$) delivery plasmid with C9 transposase, *ori pSC101* Cb$^{r}$ | (8) |
| pDL1073 | Tn*10* (Km$^{r}$) delivery plasmid with ATS transposase, *ori pSC101* Cb$^{r}$ | (9) |
| pUC18 | *oriColE1* MCS Cb$^{r}$ | (10) |
| pSR47S | *oriTRP4 oriR6K sacB* Km$^{r}$ | (11) |
| pJB4648 | Gm$^{r}$ derivative of pSR47S | (11) |
| pUC18T-miniTn7T-Gm | *oriColE1* Gm$^{r}$ Amp$^{r}$; miniTn7 base vector | (12) |
| pYDE009 | pUC18T-miniTn7T-Gm with *tetR*-tetP-*dcas9*-*rrnB*T1-T7Te fragment from pdCas9-bacteria (Addgene #44249), Gm$^{r}$, Cb$^{r}$ | This study |
| pWH1266 | *ori pBR322 ori pWH1277* MCS Tc$^{r}$ Cb$^{r}$ | (13) |
| pYDE007 | sgRNA delivery plasmid, Cb$^{r}$ (pWH1266 with $P_{J23119}$-sgRNA$_{mrfp}$-terminator), non-targeting control | This study |
| pJE53 | sgRNA delivery plasmid, Km$^{r}$ (derivative of pYDE007 with Km$^{r}$ marker replacing Cb$^{r}$) | This study |
| pJE55 | pJE53 derivative with sgRNA$_{adc}$ | This study |
| pAFE119 | pYDE007 derivative with sgRNA$_{ftsZ}$ | This study |
| pYDE317 | pYDE007 derivative with sgRNA$_{advA}$ | This study |
| pJE15 | pYDE007 derivative with sgRNA$_{RS03245}$-15 | This study |
| pJE16 | pYDE007 derivative with sgRNA$_{RS03245}$-16 | This study |
| pJE23 | pYDE007 derivative with sgRNA$_{wza\text{-}wzb\text{-}wzc}$-1 | This study |
| pJE22 | pYDE007 derivative with sgRNA$_{wza\text{-}wzb\text{-}wzc}$-2 | This study |
| pJE25 | pYDE007 derivative with sgRNA$_{wza\text{-}wzb\text{-}wzc}$-3 | This study |
| pYD318 | pYDE007 derivative with sgRNA$_{wza\text{-}wzb\text{-}wzc}$-4 | This study |
| pEGE305 | Derivative of pWH1266 *E. coli*-*A. baumannii* shuttle vector with *lacI*$^{q}$,T5-*lac*P, Tc$^{r}$ | (4) |
| pYDE152 | pEGE305 derivative with multiple cloning site polylinker, Tc$^{r}$ (identical to pYDE153) | (8) |
| pJE10 | pYDE152 with *ACX60_RS03245-gfp* (contains *lacI*$^{q}$-T5*lac*P-*ACX60_RS03245-gfp* fragment) | This study |
| pJE41 | pUC18 with *lacI*$^{q}$-T5*lac*P-*ACX60_RS03245-gfp* (conditional *RS03245* allele exchange 3’ homology arm) | This study |
| pJE40 | pUC18 with ~1kb upstream of ACX60_RS03245 (conditional *RS03245* allele exchange 5’ homology arm) | This study |
| pJE44 | pJB4648 with conditional ACX60_RS03245 allele exchange construct (3’ and 5’ homology arms), | This study |
| pJE51 | pWH1266 (*tet*$P_{con}$) with ACX60*_RS03245-gfp* | This study |
| pJE50 | pWH1266 (*tet*$P_{con}$) with ACX60*_RS03245-3XFLAG* | This study |
| pEGE76 | pSR47S containing Δ*wzc*::Gm$^{r}$ deletion construct | (5) |
