## Supplemental Table 6 for "Essential gene analysis in *Acinetobacter baumannii* by high-density transposon mutagenesis and CRISPR interference"

**Table S6. Oligonucleotide primers**

| **Cloning** | | |
|---|---|---|
| **primer name** | **Sequence (5' – 3'; restriction site underlined if present)** | **RE site(s) if present** |
| ***Primers for RS03245 deletion*** | | |
| *pJE40 construction* | | |
| AraC-upF | ACGCGTCGACTGCAGACAAACCTGCCATAA | SalI |
| AraC-upR | CATGCATGCCACTTTAAATTGACCTAAATGACCGTCTA | SphI |
| *pJE41 construction* | | |
| AraC-downF | GCTGTCAAACATGAGAATTGCTCCGACTA | |
| AraC-downR | AAATATGCGGCCGCCCGGGAGTAACACCAGTCCAC | NotI |
| ***Primers for RS03245 expression* in trans** | | |
| *pJE10 construction* | | |
| AraC-GFP-F | CGCGGATCCGTGAACATCAAAGAGACTGAAGGA | BamHI |
| AraC-GFP-R | TGCTCTAGAATGGTAACCATGGAGCTGGCGAT | XbaI |
| ***Primers for dcas9 plasmid construction*** | | |
| pYDE009-F | ACTAGTCTTAAGACCCACTTTCACATTTAAG | SpeI |
| pYDE009-R | GATACTGCAGACTTCCCTGTTAAGTATCTTCCTG | PstI |
| ***Primers for sgRNA plasmid construction*** | | |
| *pYDE007 construction* | | |
| pYDE007-F | GATCAGGATCCGATCTTTGACAGCTAGCTCAGTC | BamI |
| pYDE007-R | GATGGTCGACGGCGCTATTCAG | SalI |
| *pJE53 construction* | | |
| Kan-F | CCGGAATTCGGTTTGATTTTTAATGGATAATGTGATATAATC | EcoRI |
| Kan-R | AAAACTGCAGACAGCTTGTCTGTAAGCGGAT | PstI |
| ***Primers for amplification of sgRNA guides*** | | |
| JB-araC sgr15 | TTACACTAGTCCTGTCAAAGCCGCATTCACGAAGGTTTTAGAGCTAGAAATAGCAAG | SpeI |
| JB-araC sgr16 | TTACACTAGTAGTAACGTCTCGACATCCATGCCAGTTTTAGAGCTAGAAATAGCAAG | SpeI |
| JB-wza sgr22 | TTACACTAGTTAAACCGGAAGTAACCGCACAGCGTTTTAGAGCTAGAAATAGCAAG | SpeI |
| JB-wza sgr23 | TTACACTAGTTTAAAGCAAGAACAGAGAAAAACGTTTTAGAGCTAGAAATAGCAAG | SpeI |
| JB-wzb sgr25 | TTACACTAGTAAAAATATTCTGCCATAGGGCTAGTTTTAGAGCTAGAAATAGCAAG | SpeI |
| JB-ampC sgr41 | TTACACTAGTTTGGTGTATTGCCCGCATAAATTGGTTTTAGAGCTAGAAATAGCAAG | SpeI |
| pAFE119 sgr01 | TTACACTAGTGTTCATCTTCTATAAATTCAAATGGTTTTAGAGCTAGAAATAGCAAG | SpeI |
| pYDE317 sgr04 | TTACACTAGTTAGGCTAGCAAATAGCCCTTGTCTGTTTTAGAGCTAGAAATAGCAAG | SpeI |
| pYDE318 sgr21 | TTACACTAGTATTCTTTTAAGTCAATTGTATCTTGTTTTAGAGCTAGAAATAGCAAG | SpeI |
| SGR-R (ApaI) | AAGTGGGCCCAAGCTTCAAAAAAAG | ApaI |
| SGR-R (KpnI) | AAGTGGGCCCGGTACCAAGCTTCAAAAAAAGCACCGAC | ApaI, KpnI |
| SGR-R (BglII) | AAGTGGGCCCAGATCTAAGCTTCAAAAAAAGCACCGAC | ApaI, BglII |
